# FluoSim: simulator of single molecule dynamics for fluorescence live-cell and super-resolution imaging of membrane proteins

**DOI:** 10.1101/2020.02.06.937045

**Authors:** Matthieu Lagardère, Ingrid Chamma, Emmanuel Bouilhol, Macha Nikolski, Olivier Thoumine

## Abstract

Fluorescence live-cell and super-resolution microscopy methods have considerably advanced our understanding of the dynamics and mesoscale organization of macro-molecular complexes that drive cellular functions. However, different imaging techniques can provide quite disparate information about protein motion and organization, owing to their respective experimental ranges and limitations. To address these limitations, we present here a unified computer program that allows one to model and predict membrane protein dynamics at the ensemble and single molecule level, so as to reconcile imaging paradigms and quantitatively characterize protein behavior in complex cellular environments. FluoSim is an interactive real-time simulator of protein dynamics for live-cell imaging methods including SPT, FRAP, PAF, and FCS, and super-resolution imaging techniques such as PALM, dSTORM, and uPAINT. The software, thoroughly validated against experimental data on the canonical neurexin-neuroligin adhesion complex, integrates diffusion coefficients, binding rates, and fluorophore photo-physics to calculate in real time the distribution of thousands of independent molecules in 2D cellular geometries, providing simulated data of protein dynamics and localization directly comparable to actual experiments.

## Introduction

Critical cellular functions such as membrane adhesion, receptor-mediated signaling, or synaptic transmission, involve the diffusional trapping of specific molecules in sub-cellular compartments ^1,2^. To quantitatively describe such molecular dynamics in living cells, several fluorescence imaging techniques are currently available ^3,4^: i) single particle tracking (SPT) which resolves the motion of individual proteins at camera frame rate; ii) photo-activation and photo-bleaching methods, namely PhotoActivation of Fluorescence (PAF) and Fluorescence Recovery After Photobleaching (FRAP) which infer protein turnover at the population level; and iii) Fluorescence Correlation Spectroscopy (FCS), which analyzes molecular dynamics by correlating intensity fluctuations. More recent approaches based on single molecule localization such as PhotoActivated Localization Microscopy (PALM) ^5,6^, direct Stochastic Optical Reconstruction Microscopy (dSTORM) ^7^, and Point Accumulation In Nanoscopic Topography (PAINT) ^8^, yield images of protein distribution at improved resolution (below 50 nm), giving unprecedented information about the nanoscale organization of biological structures ^9^.

Despite such progress in imaging power, many experimental parameters remain difficult to estimate or control, including protein expression levels, probe labeling density, potential fixation artifacts, spatial and temporal sampling of the recordings, and protein motion below the system resolution, such that results from different experimental paradigms are often difficult to reconcile ^10^. Thus, there is a pressing need for computer simulators that could unify those different imaging modes in a unique framework, estimate their respective biases, and serve as a predictive tool for experimenters, with the aim to quantitatively decipher protein organization and dynamics in living cells. Several particle-based packages relying on Monte Carlo simulations already exist to predict random motion and multi-state reactions of biological molecules, but either they do not integrate fluorescence properties or are limited to a specific type of imaging mode, and are usually not performing real-time visualization ^11–18^.

In this context, we provide here fast, robust, and user-friendly software (*FluoSim*) that allows real time simulation of membrane protein dynamics in live-cell imaging (SPT, FRAP, PAF, and FCS) and super-resolution (PALM, dSTORM, uPAINT) modalities. We also show that FluoSim can be further used to produce large virtual data sets for training deep neural networks for image reconstruction ^19^. This software should thus be of great interest to a wide community specialized in imaging methods applied to cell biology and neuroscience, with the common aim to better understand membrane dynamics and organization in cells.

## Results

### General principle of FluoSim

The FluoSim interface looks like performing a real experiment: the user imports a 2D cellular geometry from a microscopy image, and populates it with a realistic number of molecules (a parameter which depends on protein expression level, cell surface area, and labeling density) **(Fig. 1a, Supplemental software and accompanying user manual)**. Kinetic parameters characterizing the diffusion and trapping of molecules in cellular regions of interest (ROI), are entered as inputs **(Supplementary Table 1)**. At each time increment (typically ~1-100 ms, adjusted to typical sensor acquisition rates), the algorithm updates the instantaneous positions of all molecules based on random number generation. Photo-switching rates of organic fluorophores or fluorescent proteins attached to the proteins of interest, further determine the fluorescence intensity associated to each molecule over time. The algorithm is optimized to visualize in real time the cellular system and provide post-processing information including molecule trajectories, image stacks, and output graphs (i.e. histograms of diffusion coefficients, FRAP, PAF, and FCS curves). Examples of simulated data sets for a realistic range of parameter values are given in **Fig. S1**. Moreover, since the positions of molecules are known with near-infinite accuracy, the program can generate super-resolved images comparable to those obtained with PALM or dSTORM, after introducing additional parameters describing protein labeling density, fluorophore duty cycle, and localization precision of the system. A complete view of the parameters used in each experimental paradigm is available from the individual examples provided in the software menu.

**Figure 1.**
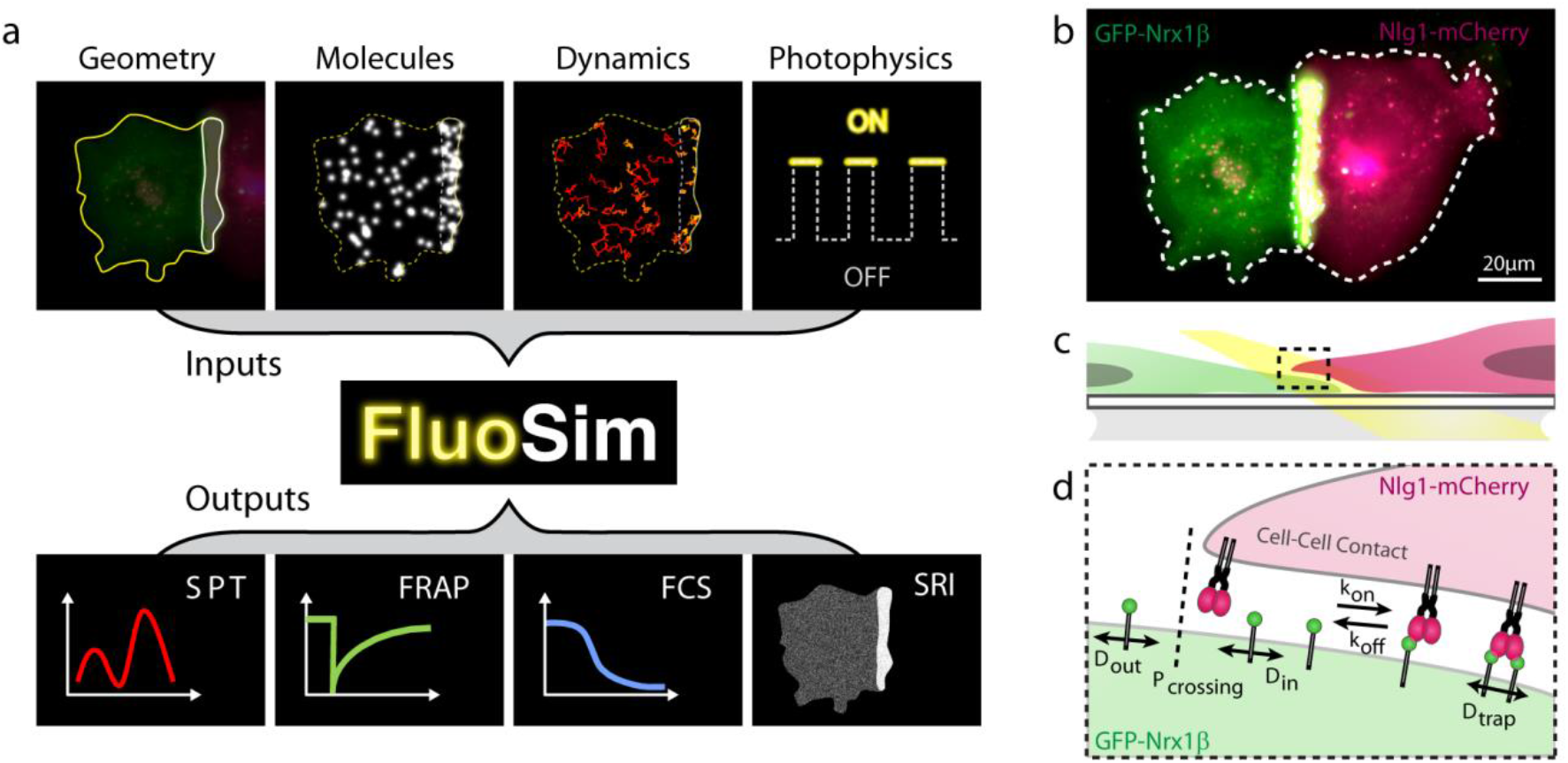
Schematics of the simulator and experimental system. **(a)** General principle of FluoSim. **(b)** Contact between two COS cells, one expressing GFP-Nrx1β (green) and the other expressing Nlg1-mCherry (magenta), resulting in molecule accumulation through adhesive interactions (yellow zone). **(c)** Diagram showing a zoomed section of the cell-cell interface at the coverslip; the yellow beam represents oblique laser illumination. **(d)** Cartoon of Nrx1β diffusional trapping by Nlg1 in a cell-cell contact.

### Experimental system to validate FluoSim

To thoroughly validate FluoSim, we performed SPT, dSTORM, FRAP, and FCS experiments essentially on the canonical neurexin-neuroligin complex that mediates trans-synaptic adhesion in neurons ^20^. We used COS-7 cells as a model expression system because they form large and flat lamellipodia that can be approximated as 2D environments for membrane diffusion. Cells were separately electroporated with recombinant GFP-tagged neurexin-1β (GFP-Nrx1β) and mCherry-tagged neuroligin-1 (Nlg1-mCherry), then cultured together for 24 h **(Fig. 1b-d)**. Both molecules reached the cell membrane and accumulated at cell-cell contacts (GFP-Nrx1β enrichment = 3.9 ± 0.5, n = 20 cells), revealing adhesive interactions.

### Simulations of SPT experiments

First, we experimentally tracked single GFP-Nrx1β molecules labeled with Atto647-conjugated anti-GFP nanobody at 50 Hz using uPAINT ^8,21,22^. Nrx1β exhibited fast free diffusion outside the adhesive contact (D_out_ = 0.3 μm²/s), and was slowed down by ~10-fold in the contact, reflecting the formation of Nrx1β-Nlg1 bonds between apposed membranes (D_trap_ =μm²/s) **(Fig. 2a,c)**. Nrx1β molecules often bounced at the contact border, revealing steric hindrance to penetrate the narrow cell-cell junction ^23^, an effect which was described by a crossing probability (P_crossing_ < 1) **(Fig. 2d)**. Next, we simulated Nrx1β diffusion using FluoSim **(Fig. 2b)**. We used the diffusion coefficients obtained experimentally and defined a sparse number of molecules (250) corresponding to the number of experimental detections per frame, together with two kinetic rates describing the Nrx1β-Nlg1 interaction taken from the literature (k_on_ = 0.15 s^−1^ and k_off_ = 0.015 s^−1^) ^24,25^. Using these parameters, FluoSim generated diffusion coefficient distribution curves inside and outside the contact that aligned quite well on experimental data **(Fig. 2c)**. Experimental distributions were somewhat more spread than theoretical ones, most likely because of local membrane heterogeneities which are not accounted for in the model.

**Figure 2.**
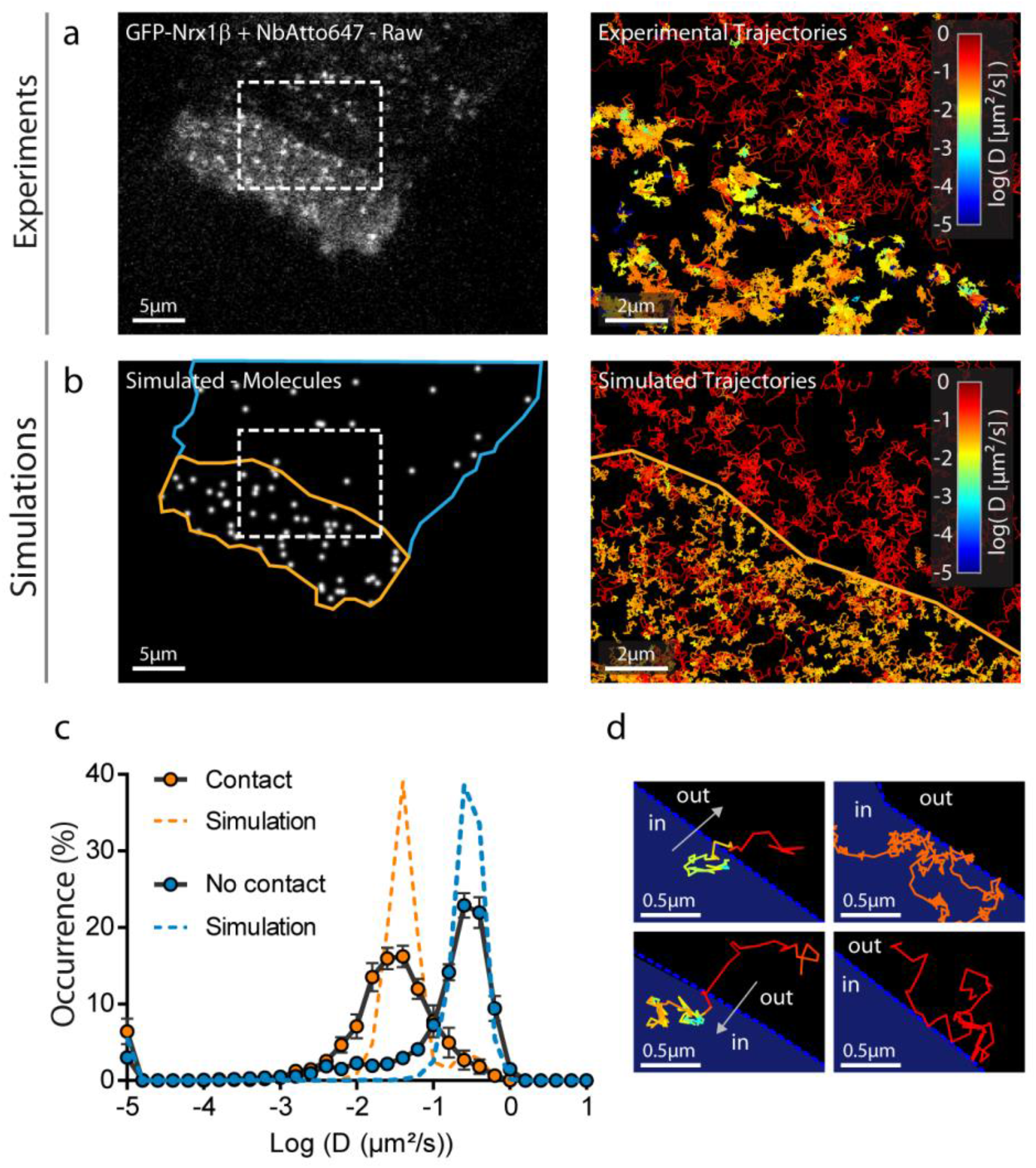
Fitting SPT experiments. **(a)** Raw image of a COS-7 cell expressing GFP-Nrx1β sparsely labeled with Atto647-conjugated GFP nanobody (white signal). **(b)** Image of simulated molecules in the same geometry. Rectangles highlight the interface between the cellular region with freely diffusing GFP-Nrx1β molecules (blue outline), and the contact region with a cell expressing Nlg1-mcherry (yellow outline). On the right, a zoom on this ROI shows single molecule trajectories for both experiments and simulations. The diffusion coefficient expressed in log scale is color coded. **(c)** Distributions of GFP-Nrx1β diffusion coefficients for experiments (circles, average ± sem of 3 cells, 4145 trajectories in contact and 4867 outside contacts) and simulations (dashed lines, 5 repetitions, 8757 trajectories in contact and 2645 trajectories outside contacts) on a semi-log plot. The Spearman correlation coefficient comparing experiment and simulation was r = 0.85 (P< 0.001, n = 36 bins) for contact regions, and r = 0.81 (P < 0.001, n =36 bins) for outside regions. **(d)** Representative examples of single molecule trajectories at the interface: (left) a molecule escapes the contact and diffuses out freely, or enters the contact and gets trapped; (right) a molecule stays stuck in the contact with low diffusion, or bounces on the contact without entering it.

### Simulations of dSTORM experiments

To simulate dSTORM experiments that were experimentally performed on GFP-Nrx1β using saturating labeling with Alexa647-conjugated GFP nanobody **(Fig. 3a)**, we defined a relatively large number of molecules in the imported geometry (70,000, corresponding to the sum of experimental detections per frame integrated over the cell surface area), and set all diffusion coefficients to zero to mimic cell fixation. To simulate stochastic fluorescence emission of Alexa 647 ^26^, we calculated the switch-on (0.006 s^−1^) and switch-off (9.3 s^−1^) rates of Alexa 647 fluorophores from isolated Alexa 647-conjugated nanobodies in dSTORM imaging conditions **(Fig. 3c)**. We then simulated the accumulation of single molecule localizations for 80,000 frames to mimic the experimental super-resolved maps of Nrx1β distribution **(Fig. 3b)**.

**Figure 3.**
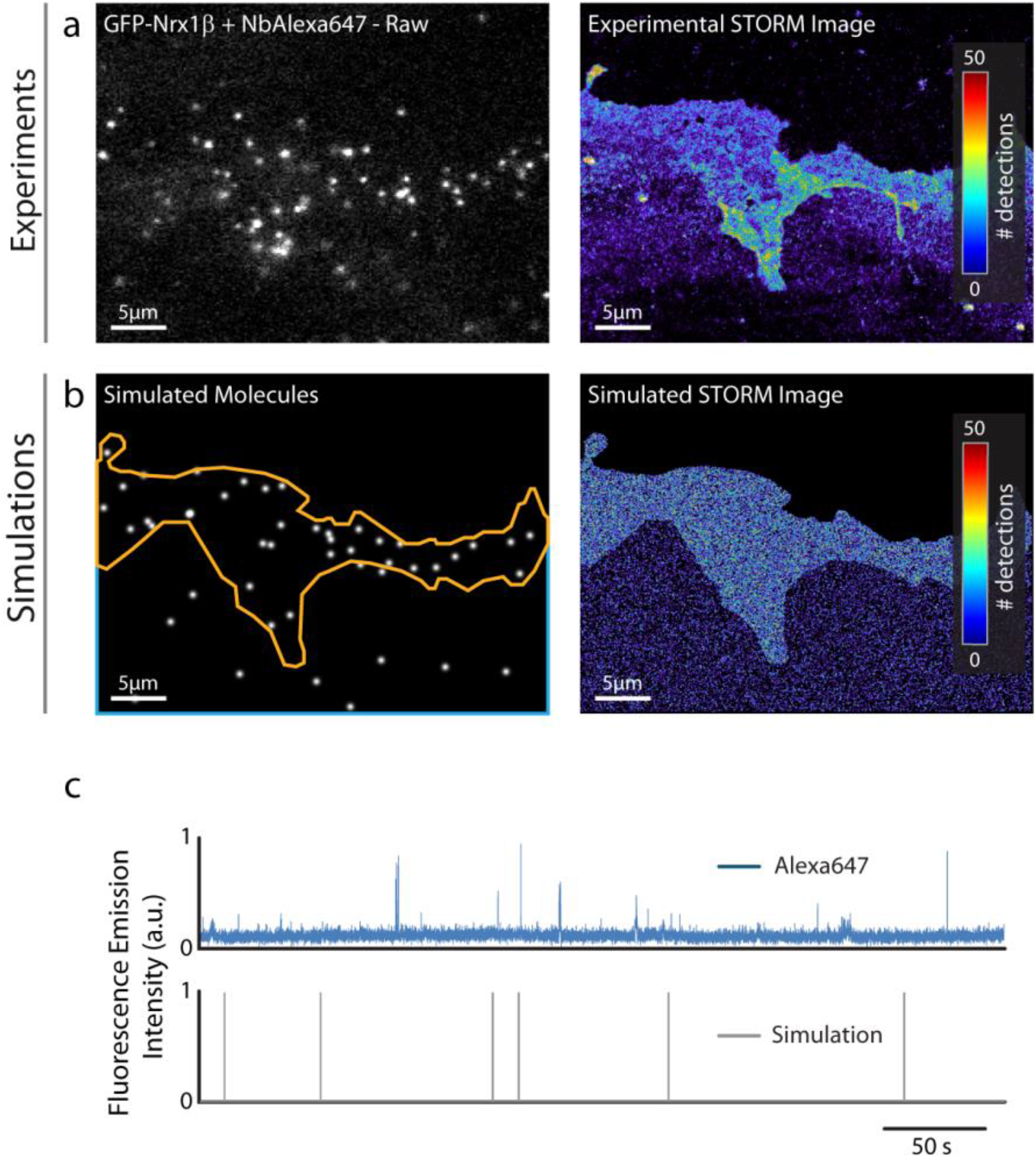
Fitting SRI experiments. **(a)** Representative single frame image of a STORM sequence performed on GFP-Nrx1β labeled with Alexa647-conjugated GFP nanobody, and corresponding super-resolved image generated from 3.57 × 10^6^ single molecule localizations (pixel size 32 nm, total acquisition time 1,600 s). The number of molecules detected per pixel is color coded. **(b)** Simulated image showing single molecule fluorescence emission in the same cell geometry, and corresponding super-resolved map with a localization precision of 58 nm (FWHM). The total number of detections is 3.61 × 10^6^. **(c)** Fluorescence emission over time from an isolated Alexa647-conjugated GFP nanobody bound to the glass coverslip in a STORM sequence, and simulated emission of fluorescence of an immobile molecule obtained with switch-on and -off rates of s^−1^ and 9.3 s^−1^, respectively.

### Training of deep neural networks for image reconstruction

Deep learning is becoming increasingly popular for image reconstruction in fluorescence microscopy ^19,27,28^. Convolutional Neural Networks (CNNs) are especially relevant for image treatment and have to be trained using large exemplary data sets obtained either experimentally, or from simulations. In this context, we tested the ability of FluoSim to train deep CNNs based on simulated data. We generated large image sets of randomly distributed single molecules represented as Gaussian intensity profiles plus Poisson noise, together with their localization maps as ground truth, and trained the previously described CNN called Deep-STORM ^19^. FluoSim-trained Deep-STORM was able to faithfully reconstruct super-resolved maps of both simulated and experimentally-observed microtubules, from single molecule localization stacks **(Figs. S2-S3)**. Strikingly, the CNN worked well at both low and high single molecule density, thus offering a considerable gain in acquisition time (×20) for an equivalent resolution ^27^. FluoSim-trained Deep-STORM also allowed the reconstruction of images of Nrx1β-Nlg1 contacts that were comparable to images obtained with PALM-Tracer ^29^, or to images directly simulated by FluoSim **(Fig. S4)**. FluoSim can therefore be used to train deep CNNs.

### Simulations of FRAP experiments

To challenge the simulator against ensemble measurements, we performed FRAP experiments on GFP-Nrx1β expressed in COS-7 cells. GFP-Nrx1β accumulates at cell-cell contacts when the opposite cell expresses its molecular partner Nlg1 **(Fig. 4a)** and shows slower recovery in the adhesive contact as compared to membrane regions not in contact with other cells **(Fig. 4b,e)**. To mimic these experiments, we introduced a large number of molecules in the simulator (up to 150,000) and generated fluorescence-like images by defining a Gaussian intensity profile for each GFP-tagged molecule (FWHM = 0.17 μm). Taking kinetic parameters from the literature (k_on_ = 0.15 s^−1^ and k_off_ = 0.015 s^−1^) and an intermediate porosity (P = 0.3), the simulated images at steady state predicted Nrx1β enrichment in the contact area that matched experimental values **(Fig. 4c,d)**. To induce local photo-bleaching, we chose a bleaching rate (4.25 s^−1^) reproducing the initial drop of fluorescence observed experimentally (~75% in 400 ms). Using those coefficients plus the diffusion parameters determined from SPT, FRAP simulations accurately reproduced experimental data **(Fig. 4e)**.

**Figure 4.**
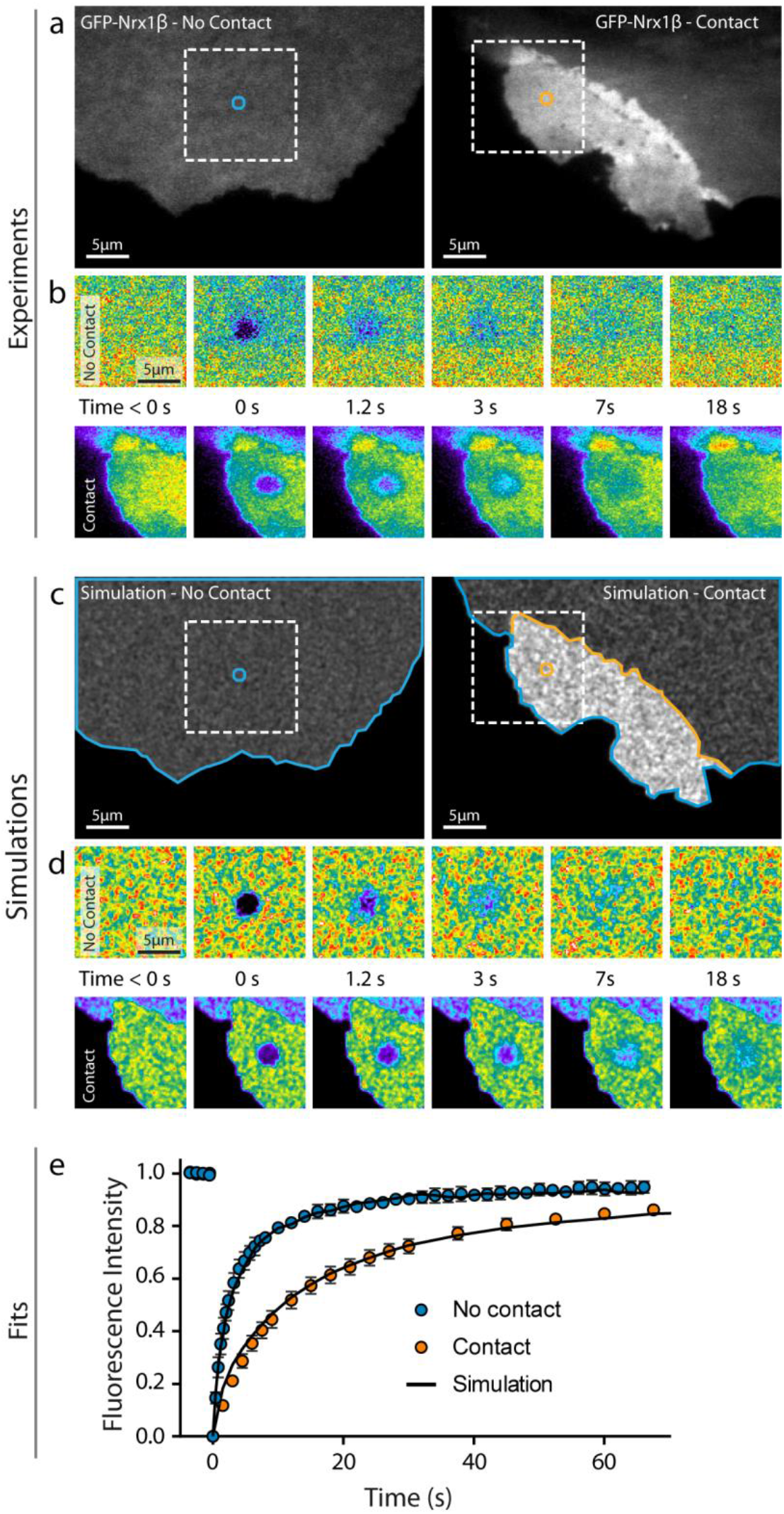
Fitting FRAP experiments. **(a)** Representative images of COS cells expressing GFP-Nrx1β either not forming contact (left), or forming contact with another cell expressing Nlg1-mCherry (right). **(b)** Corresponding FRAP sequences on the zoomed square areas (bleached circle diameter = 2.8 μm). Intensity is color coded. **(c, d)** Simulated images of the same cell geometries filled with 150,000 fluorescent molecules, and corresponding FRAP sequences. **(e)** Normalized FRAP curves obtained by experiment (dashed) outside (mean ± sem of 6 cells, 44 bleached regions) or inside contact regions (mean ± sem of 18 cells, 144 bleached regions) and corresponding simulations (solid curves, average of 30 repetitions each, sem < 1% mean, not shown). The Spearman correlation coefficient comparing experiment and simulation was r = 1.0 (P< 0.001, n = 21 time points) for contact regions, and r = 0.98 (P < 0.001, n =43 time points) for outside regions.

### Simulations of FCS experiments

To further validate the simulator in conditions of intermediate molecular density, we performed FCS experiments ^30^ by recording intensity fluctuations of Atto647-nanobody in a diffraction-limited laser spot at 200 Hz **(Fig. 5a,b)**. As expected from slower diffusion, the autocorrelation function obtained for Nrx1β in the adhesive contact was shifted to the right compared to the free region, with a small photo-bleaching bias due to longer residence times **(Fig. 5e)**. Again, the simulations performed with an intermediary number of molecules introduced in FluoSim, matched experimental FCS curves with the same set of kinetic and photophysical parameters used for SPT and FRAP **(Fig. 5c-e)**.

**Figure 5.**
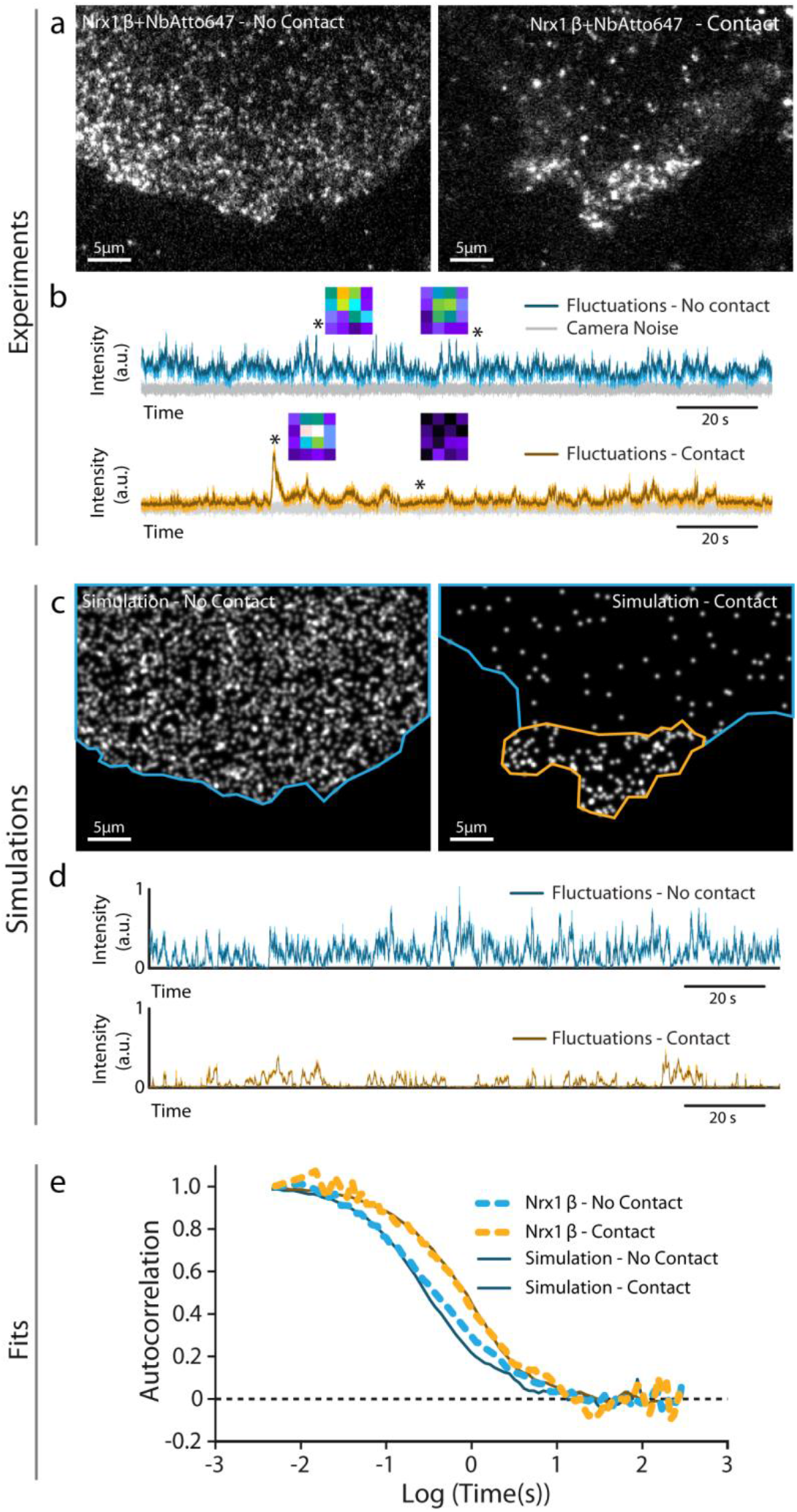
Fitting FCS experiments. **(a)** Images of Atto647-conjugated GFP nanobody bound to COS cells expressing GFP-Nrx1β not forming contact (left), or forming contact with a cell expressing Nlg1-mCherry (right). A 642 nm focused laser beam of 0.6 μm FWHM was parked in contact or no-contact region, and square images of the illuminated area (1 μm × 1 μm) were collected at 200 Hz. **(b)** Intensity fluctuations over time in the two regions (color), above the camera noise (grey). Insets represent images acquired at the indicated times (stars). Intensity is color coded. **(c, d)** Simulated images using the same cell geometries populated with 2500 and 200 molecules, respectively, and corresponding intensity fluctuations. **(e)** Normalized autocorrelation functions for FCS experiments (dashed) performed outside (2 cells / 10 recordings) and inside contact regions (1 cell / 3 recordings), and corresponding simulations (solid, average of 10 repetitions for each curve).

## Discussion

In summary, FluoSim allows a prediction and comparison of membrane protein dynamics in a wide range of fluorescence cell imaging modes, with a precise control of the relevant kinetic, photo-physical, and acquisition parameters determining measurement outputs. The program is intended to help biologists adjust and interpret their experiments on a variety of cellular systems, and serve as a teaching resource in bio-imaging programs.

FluoSim reproduced a wide range of experimental results (SPT, FRAP, FCS, SRI) on the Nrx1β-Nlg1 membrane complex using a unique set of parameters extracted from published in-vitro studies and/or taken from our own measurements ^24,25^, thereby giving strong credit to the correlative approach. The program is very fast and robust, and should be applicable to model a wide range of 2D-like dynamic molecular systems experiencing membrane diffusion and transient confinement, for example integrins at focal contacts in fibroblasts ^31,32^, cadherins at cell-cell contacts ^33–35^, neuronal adhesion proteins and neurotransmitter receptors at synapses ^22,36,37^, and trapping of membrane molecules by lipid raft molecules or cytoskeletal interactions ^38–40^.

We believe that this quantitative simulation approach will be of great help to optimize experimental design, especially regarding the choice of various parameters such as fluorophores, control of laser powers, acquisition frame rate, and overall timing of the experiment with respect to the internal dynamics of the molecular system, thus replacing lengthy experimental adjustments and saving research time. The FluoSim code might even be integrated into existing microscopy packages to provide a feedback loop to the image acquisition parameters. The software can also be used to test the robustness and predictions of single molecule tracking algorithms that have been implemented those past years, e.g. SR-Tesseler and InferenceMap ^41,42^. Compared to existing software such as PyFRAP, SuReSim, FERNET, or MCell ^13–15^, FluoSim integrates many fluorescence modalities into a single program and achieves real-time display **(Supplemental Table 2)**. In its present version, FluoSim is limited to Brownian motion and first order molecular reactions, but sub-or super-diffusive behaviors as well as more complex multi-state molecular reactions might be implemented on a case-by-case basis, depending on user needs.

Finally, with the advent of single-molecule based super-resolution microscopy such as PALM and STORM, many new questions arise regarding the degree of labeling needed, and the number of single molecule localizations to accumulate in order to reconstruct a realistic image of biological structures, with the risk of finding artificial properties in under-sampling conditions ^10^. By varying the number of molecules, type of fluorophore attached to them, localization precision, and length of the acquisition sequence, the software is able to determine the conditions for faithful detection of the biological sample. The capacity of FluoSim to mimic molecule dynamics might also be used in the near future to train the next generation of CNNs for the analysis of live imaging experiments ^43^.

## Methods

### DNA plasmids

GFP-Nrx1β was a kind gift from M. Missler (Münster University, Germany). HA-tagged Nlg1 obtained from P. Scheiffele (Biozentrum, Basel) was used as a backbone to construct NLG1-mCherry, by inserting mCherry intracellularly at position -21aa before the C-terminus. The HA-NLG1 sequence was moved from the pNice vector into a pcDNA vector at the HindIII/ NotI sites. Two PCRs were performed on Nlg1: one from KpnI (inside the Nlg1 sequence) at the insert position of mCherry (AgeI site added) and one of the end of Nlg1 (NheI / NotI). The mCherry gene (AgeI / NheI) was obtained by PCR on pmCherry-N1 (Clontech). A four-fragment-ligation was then done to obtain the final construct (HindIII-HA-Nlg1-AgeI-mCherry-NheI-Nlg1Cter-NotI). The plasmid for bacterial expression of the anti-GFP nanobody ^44^ was a kind gift from A. Gautreau (Gif-sur-Yvette, France). The bacterial production of anti-GFP nanobody, purification, and conjugation to organic dyes (Atto647N or Alexa 647), was described previously ^22^.

### Cell culture and electroporation

COS-7 cells (ATCC) were cultured in Dulbecco’s modified Eagle’s medium (DMEM; GIBCO/BRL) supplemented with 10% fetal bovine serum (FBS), 100 units mL^−1^ penicillin, and 100 μg mL^−1^ streptomycin, in a 37°C-5% CO_2_ atmosphere. One day before the experiments, cells were rinsed twice in warm PBS, trypsinized for 5 min, mixed in culture medium, and centrifuged for 5 min at 1,000 rpm. The cell pellet was suspended in 100 μL electroporation medium and electroporated for either GFP-Nrx1β or NLG1-mCherry plasmids with the Amaxa Nucleofector system (Lonza), using 500,000 cells per cuvette and 3 μg DNA. Electroporated cells were mixed in culture medium, seeded on 18 mm glass coverslips at a concentration of 50,000-80,000 cells per coverslip, cultured in 12-well plates, and imaged 24-48 hrs after electroporation.

### Single Molecule Tracking (uPAINT)

Universal point accumulation in nanoscale topography (uPAINT) experiments were carried out as reported ^8^. Cells were mounted in Tyrode solution (15 mM D-glucose, 108 mM NaCl, 5 mM KCl, 2 mM MgCl_2_, 2 mM CaCl_2_ and 25 mM HEPES, pH 7.4) containing 1% globulin-free BSA (Sigma A7638) in an open Inox observation chamber (Life Imaging Services, Basel, Switzerland). The chamber was placed on a fully motorized inverted microscope (Nikon Ti-E Eclipse) equipped with perfect focus system, a thermostatic box (Life Imaging Services) providing air at 37°C, and an APO TIRF 100x/1.49 NA oil immersion objective. GFP-and mCherry expressing cells were detected using a mercury lamp (Nikon Xcite) and the following filter sets (SemROCK): EGFP (Excitation: FF01-472/30; Dichroic: FF-495Di02; Emission: FF01-525/30) and mCherry (Excitation: FF01-543/22; Dichroic: FF-562Di02; Emission: FF01-593/40). Cells expressing GFP-Nrx1β were labeled using a low concentration of Atto647N-conjugated GFP nanobody (1 nM). A four-colour laser bench (405/488/561nm lines, 100 mW each; Roper Scientific, and 1 W 647 nm line, MPB Communications Inc., Canada) is connected through an optical fiber to the Total Internal Reflection Fluorescence (TIRF) illumination arm of the microscope. Laser power was controlled through an acousto-optical tunable filter (AOTF) driven by the Metamorph^®^ software (Molecular Devices). Atto 647N was excited with the 647 nm laser line (~2 mW at the objective front lens), through a four-band beam splitter (BS R405/488/561/635, SemRock). Samples were imaged by oblique laser illumination, allowing the excitation of individual Atto-conjugated ligands bound to the cell surface, without illuminating ligands in solution. Fluorescence was collected on an EMCCD camera with 16 μm pixel size (Evolve, Roper Scientific, Evry, France), using a FF01-676/29 nm emission filter (SemRock). Stacks of 4,000 consecutive frames were obtained from each cell with an integration time of 20 ms, using the Nikon perfect focus system to avoid axial drift. Images were analyzed using PALM-Tracer, a program running on Metamorph^®^ and based on wavelet segmentation for molecule localization and simulated annealing algorithms for tracking (generously provided by JB Sibarita, Bordeaux) ^45^. This program allows the tracking of localized molecules through successive images. Trajectories longer than 20 frames (400 ms) were selected. The diffusion coefficient, D, was calculated for each trajectory, from linear fits of the first 4 points of the mean square displacement (MSD) function versus time. Trajectories with displacement inferior to the pointing accuracy (~50 nm in uPAINT conditions) whose MSD function cannot be fitted are arbitrarily taken as D = 10^−5^ μm² s^−1^.

### dSTORM experiments

COS-7 cells expressing GFP-Nrx1β were surface-labeled with a high concentration (100 nM) of Alexa647-conjugated GFP Nanobody in Tyrode solution containing 1% globulin-free BSA (Sigma A7638) for 10 min, rinsed and fixed with 4% PFA-0.2% glutaraldehyde in PBS for 10 min at room temperature, and stored in PBS at 4°C until imaging. For microtubule staining, COS-7 cells were rinsed twice in PBS, fixed using 4% PFA-20% sucrose for 15 min at room temperature, washed 3 times in PBS, and incubated with 50 mM NH_4_Cl in PBS for 10 min. After fixation, cells were washed 3 times in PBS, permeabilized using 0.2% Triton-X 100 for 10 min, washed in PBS, blocked in 1% BSA-PBS for 30 minutes, and incubated with anti-α-tubulin (Thermofisher MA1-80017, 1/500) overnight at 4°C. The next day, cells were washed 3 times in PBS, and incubated 45 min with secondary goat-anti-rat-Alexa647 antibody (ThermoFisher A21247, 1/800) and kept in PBS before dSTORM imaging.

Cells were imaged in Tris-HCl buffer (pH 7.5), containing 10% glycerol, 10% glucose, 0.5 mg/mL glucose oxidase (Sigma), 40 mg/mL catalase (Sigma C100-0,1% w/v) and 50 mM β-mercaptoethylamine (MEA) (Sigma M6500) ^46^. The same microscope described for uPAINT was used. Pumping of Alexa647 dyes into their triplet state was performed for several seconds using ~60 mW of the 647 nm laser at the objective front lens. Then, a lower power (~30 mW) was applied to detect the stochastic emission of single-molecule fluorescence, which was collected using the same optics and detector as described above for uPAINT. 10-20 streams of 3,000-4,000 frames each were acquired at 50 Hz. To generate images intended to test deep CNN algorithms, a higher density of fluorescent molecules was generated by turning on the 405 nm laser power to 5 mW during acquisition. Multi-color 100 -nm fluorescent beads (Tetraspeck, Invitrogen) were used to register long-term acquisitions and correct for lateral drift. The localization precision of our imaging system in STORM conditions is around 60 nm (FWHM). Stacks were analyzed using the PALM-Tracer program, allowing the reconstruction of a unique super-resolved image of 32 nm pixel size (zoom 5 compared to the original images) by summing the intensities of all localized single molecules (1 detection per frame is coded by an intensity value of 1).

### FRAP experiments and analysis

COS-7 cells expressing GFP-Nrx1β in co-culture with cells expressing NLG1-mCherry were mounted in Tyrode solution, and observed under the same set-up used for uPAINT and STORM. The laser bench has a second optical fiber output connected to an illumination device containing two x/y galvanometric scanning mirrors (ILAS, Roper Instrument) steered by MetaMorph. It allows precise spatial and temporal control of the focused laser beam at any user-selected region of interest within the sample for targeted photo-bleaching. Switching between the two fibers for alternating between imaging and bleaching is performed in the ms time range using an AOTF. Oblique illumination acquisition was performed using the 491 nm at low power (0.3 mW at the front of the objective) to image molecules in the plasma membrane close to the substrate plane. After acquiring a 10 sec baseline at 1 Hz frame rate, rapid selective photo-bleaching of 3-9 circular areas of diameter 2.8 μm was achieved at higher laser power (3 mW at the objective front lens), during 400 ms. Fluorescence recovery was then recorded immediately after the bleach sequence for 80 sec. The recording included three phases with decreasing frame rate ranging from 2 to 0.1 Hz. Observational photo-bleaching was kept very low, as assessed by observing control unbleached areas nearby. FRAP curves were obtained by computing the average intensity in the photobleached area, after background subtraction, and normalized between 1 (baseline) and 0 (time zero after photo-bleaching).

### FCS experiments and analysis

COS-7 cells expressing GFP-Nrx1β were surface-labeled at an intermediate concentration (10 nM) of Atto647-conjugated GFP nanobody for 5 min at 37°C, then mounted in Tyrode solution on the same microscope. The 642 nm laser beam was parked on a region of interest, either on the free cell membrane, or in the cell-cell contact area. The laser was kept at low power (3% of 100 mW, i.e. ~0.2 mW at the objective front lens) to avoid photo-bleaching the organic dye. Using glass coverslips coated with polylysine and higher concentrations of Atto647-conjugated nanobody (100 nM) to provide a uniform density, we independently measured the Gaussian intensity profile of the laser beam, which gave a standard deviation σ = 0.26 μm (FWHM = 0.6 μm). Intensity fluctuations due to Nrx1β molecules entering and leaving the laser spot by membrane diffusion, were detected using a second camera (Hamamatsu Orca Flash 4.0) on the opposite port of the microscope. A square area of 16 × 16 pixels (1 μm × 1μm) centered on the laser spot was chosen, and streams of 60,000 images at binning 4 were acquired through the Hokawo software (Hamamatsu) at 200 Hz or 500 Hz, for the study of cell-cell contacts or free regions, respectively. Control stacks with the laser off were acquired to record the camera noise. The integrated intensity of each image was read from the multi-TIFF stacks, from which the average camera noise was subtracted, and the autocorrelation function was computed using custom-made routines written in C++, and normalized by its first value.

### Description of the simulator

#### Programing details

The FluoSim source code has been written with the C++ programming language using the ISO/IEC 14882:2011 standard (C++11 standard). The FluoSim program and its related libraries have been compiled with the 32bit MinGW compiler (MinGW 4.8 32bit) and can be executed on 32-bit and 64-bit Windows operating systems. The FluoSim project has been developed using the Integrated Development Environment QtCreator 3.0.1.

#### Libraries

The C++ standard library has been extensively used to write the FluoSim source code. The Qt library (Qt 5.2.1) has been used to implement the Graphical User Interface. To allow live rendering of the simulations, FluoSim benefits from hardware acceleration through the openGL library (openGL 3.2). OpenGL extensions have been loaded with the glew library. Vectors and matrices manipulations utilized in FluoSim calculations are implemented in the glm library. The lmfit library which contains functions to perform non-linear fitting has been used during MSD fitting. Qt, openGL, glew and glm libraries are dynamically linked to FluoSim while the lmfit library has been directly integrated in the source code. The program is provided as an executable file within a folder containing the necessary .dll files to operate properly (see the user manual for a description of how to run the program).

#### Code availability

The software is freely available for academic use as Supplementary Software or upon request to the corresponding author. Source code will be made available to the scientific community after manuscript acceptance.

#### General algorithm

Our computational approach is based on previously reported frameworks to describe AMPA receptor trafficking at synapses ^47^ and actin retrograde flow in growth cones^34^. However, whereas in previous programs the simulations were run one by one, and later visualized using a commercial image analysis software (Metamorph, Molecular Devices), FluoSim is a stand-alone program that allows the fast calculation of thousands of single molecule positions in parallel, compatible with live image rendering. A further important improvement over previous approaches is that the working space is now determined from an imported microscopy image with potentially complex shapes. The cell outline is imported as a region file previously made in Metamorph or Image J, or directly drawn on the screen using a toolbox. This internal space is randomly populated by a given number of molecules (1-200,000), GFP-Nrx1β in our case. Those molecules are kept within the cell boundaries by rebound conditions. An individual molecule is characterized by its 2D coordinates × and y over time t, and its intensity. The total duration of the simulations (typically 5 s-10 min) is set according to the experiment to model. The time step of the simulations *Δt* is varied between 1-100 ms, corresponding to typical detector frame rates in FCS and SPT experiments, respectively. The initial position of a freely diffusing molecule is defined by x(0) = x_0_ and y(0) = y_0_, taken as random numbers to fall within the cell boundaries. The diffusion coefficient outside the contact area (*D*_*out*_) is chosen around 0.3 μm²/s, based on SPT data, while the contact area can be characterized by a lower diffusion coefficient (*D*_*in*_ in the range of 0.1-0.3 μm²/s), owing to molecular crowding and steric hindrance. An additional coefficient called crossing probability (*P*_*crossing*_ between 0 and 1) describes the potentially limited penetrability of molecules into the contact. A small fraction of immobile Nrx1β molecules was observed (~5% with D < 10^−3^ μm²/s), which might be due to non-specific adhesion or endocytosis^48^, and introduced in the program at random positions with zero diffusion coefficient. In the contact area, surface-diffusing Nrx1β and Nlg1 molecules are allowed to bind reversibly, with first order binding and unbinding rates *k*_*on*_ and *k*_*off*_, respectively (both in s^−1^). The *k*_*off*_ value was taken from surface plasmon resonance data obtained on purified extracellular domains of Nrx1β and Nlg1 ^24^ (0.015 s^−1^), while *k*_*on*_ was inferred from previous experiments of Nrx1β-coated Quantum dots interacting with neurons expressing Nlg1 ^25^ (0.15 s^−1^). Bound complexes were allowed to diffuse at a lower diffusion coefficient *D_trap_* = 0.04 μm²/s, reflecting their slow movement within the cell-cell contact. The number of Nlg1-mCherry binding sites is assumed to be in excess (consistently with high expression levels in COS-7 cells), such that the binding rate *k*_*on*_ is maintained constant throughout the simulations, i.e. it does not depend on the number of Nrx1β-Nlg1 complexes formed over time. We further consider a uniform (not a discrete) distribution of binding sites in the trapping area, also consistent with a high density of Nlg1 molecules.

#### Calculation of positions

At each time step, the (x,y) coordinates of each molecule are incremented by the distances (Δx, Δy), which depend on whether the molecule is outside or inside the contact area, or in an adhesive complex. If the molecule is outside the contact area, it follows a random walk with diffusion coefficient *D*_*out*_. The positions x(t) and y(t) are then incremented at each time step by n_x_(2*DΔt*)^1/2^ and n_y_(2*DΔt*)^1/2^, respectively, where nx and ny are random numbers generated from a normal distribution, to account for the stochastic nature of diffusion. This ensures that the mean square displacement stays proportional to time, i.e. <x^2^ + y^2^> = 4*D_out_t*. If the adhesion molecule reaches a contact area, it is set to diffuse with a lower diffusion coefficient *D_in_*, with increments n_x_(2*D_in_Δt*)^1/2^ and n_y_(2*D_in_Δt*)^1/2^. Whenever the molecule reaches the contact area, it is allowed to bind to its counter-receptor only if the probability of coupling in this time interval, *P*_*coupl*_ = *k*_*on*_*Δt*, is greater than a random number *N* between 0 and 1 generated from a uniform distribution. If this is not the case, the molecule continues to diffuse until both conditions are met, i.e. it is within the contact area and the probability of binding is greater than the random number *N*, chosen different at each time increment. Upon binding, the adhesive complex is set to diffuse with a slow diffusion coefficient *D_trap_*, thus the positions x(t) and y(t) are incremented by n_x_(2*D*_*trap*_*Δt*)^1/2^ and n_y_(2*D*_*trap*_*Δt*)^1/2^, respectively. The complex stays bound until the probability for dissociation *P*_*detach*_ = *k*_*off*_*Δt*, exceeds another random number *N’*. It then binds again or escapes into the contact or outside space. Starting with random positions, it can take a relatively long time before molecules reach a steady-state distribution. Yet, it is necessary that the molecular system is at steady-state before recording a given simulation. To accelerate this process, an option is proposed in FluoSim to theoretically estimate the steady-state, by placing more molecules in the membrane compartments, considering both slower diffusion and adhesion. The molecular enrichment was then given by the formula (*P*_**crossing**_ *D*_*out*_ /*D*_*in*_) (1+*k*_*on*_/*k*_*off*_). Using those dynamic coefficients, we chose the value of *P*_*crossing*_ for each type of simulation (0.25-0.7), to match the experimental enrichment of GFP-Nrx1β in the cell contact normalized to outside areas.

#### Photophysics and molecule intensity

In addition to its position, each molecule is defined by its fluorescence intensity over time which can be either 0 or 1. Intensity is set by two photo-physical parameters: the switch-on rate (*k*_*on*_^*Fluo*^) and the switch-off rate (*k*_*off*_^*Fluo*^). These rates are in units of sec and represent the probabilities per unit of time that a molecule will switch from a state where it emits fluorescence, to a state where it does not emit fluorescence, and vice versa. The rates are specific for each fluorophore (GFP, mCherry, mEos2, Atto dyes) and strongly depend on the laser powers used to image them. By playing on these two rates, many types of experiments can be mimicked. For example, to model a PALM or STORM experiment, one sets a low switch-on rate to induce sparse stochastic emission and a high switch-off rate to induce rapid extinction of fluorescence. In a PAINT experiment, *k*_*on*_^*Fluo*^ represents instead the rate of binding of fluorescent ligands in solution to receptors on the cell surface, which spontaneously appear in the oblique illumination plane, whereas *k*_*off*_^*Fluo*^ combines fluorophore photo-bleaching and probe detachment from the cell surface. To mimic a FRAP experiment, *k*_*off*_^*Fluo*^ is set to a high level in a given ROI to quickly and irreversibly photo-bleach fluorophores, then monitor recovery. Conversely, in a Photo-Activation of Fluorescence (PAF) experiment, *k*_*on*_^*Fluo*^ is set to a high value in the ROI to be activated, and the fluorescence redistribution is followed. In FCS experiments, a low value of *k*_*off*_^*Fluo*^ can be introduced to reproduce observational photo-bleaching.

#### Running simulations and data export

Once realistic parameters have been tested in live mode, simulations can be generated, and results are exported in various forms. For SPT simulations,.trc files containing the spatial positions and intensity of each molecule over time are saved, and can be loaded later for offline visualization and analysis (menu SPT Analysis). For FRAP, PAF, and FLIP simulations, average intensities over time in defined ROIs are saved as txt files. For FCS simulations, both intensity fluctuations and autocorrelation values over time are exported as .txt files. For SRI simulations, a single super-resolved image integrating all single molecule localizations is exported as a TIFF file.

### SPT simulations

To mimic the sparse density of GFP-Nrx1β bound to Nanobody-Atto647 as used in uPAINT experiments, a low number of molecules were introduced in the model cell (250 molecules corresponding to a surface density of 0.43 mol/μm²). The length of the simulated trajectories was adjusted to the experimental one by choosing *k*_*off*_^*Fluo*^ = 3 s^−1^ corresponding to a mean trajectory duration of 340 ms (i.e. 17 frames of 20 ms each). The parameter *k*_*on*_^*Fluo*^ which determines the number of fluorescent molecules was set to 1 s^−1^, so as to yield approximately the same density of visible molecules per surface area as in the experiments (0.1 mol/μm²). Sequences of 4,000 frames were generated as in the experiments, and only trajectories longer than 20 frames were selected (total 2244 trajectories). The diffusion coefficient, *D*, was calculated for each trajectory, from linear fits of the first 4 points of the MSD function versus time. Five independent simulations were run for each set of parameters, allowing the construction of histograms of diffusion coefficients directly comparable to SPT experiments.

### FRAP simulations

To match the very dense distribution of GFP-Nrx1β molecules that characterize FRAP experiments, a large number of molecules were introduced in the virtual cell (150’000 molecules corresponding to a surface density of ~200 mol/μm²). Simulations of 1,000 frames, including a baseline of 100 frames, were generated with a time step of 100 ms (total duration 100 s). Two areas were recorded, one within the adhesive contact, the other outside (bleached diameter = 2.8 μm). The photoactivation rate was set to a maximal value (*k*_*on*_^*Fluo*^ = 5 s^−1^), i.e. all molecules are initially fluorescent, while the photo-bleaching rate is set to zero during baseline and recovery acquisition (i.e. observational photo-bleaching is neglected here). During the short photo-bleaching period (400 ms), the photo-bleaching rate is set to *k*_*off*_^*Bleach*^ = 4.25 s^−1^ for 4 frames, to precisely match the initial drop of fluorescence observed experimentally (~75%). The number of molecules in the photo-bleached areas was computed over time, and normalized between 1 (baseline number of fluorescent molecules before photo-bleaching) and zero (number of fluorescent molecules right after photobleaching). FRAP simulations were repeated 10 times, and the corresponding curves were averaged.

### FCS simulations

To mimic the intermediate densities of GFP-Nrx1β bound to Nanobody-Atto647 used in FCS experiments, 200 or 2500 molecules (corresponding to 0.27 and 2.8 mol/μm²) were entered in the program, to simulate cells forming or not forming adhesive contacts, respectively. These values roughly correspond to the experimental labeling densities, and ensure large enough fluctuations to calculate a reliable autocorrelation function. Simulations of 500,000 or 1,000,000 frames with time steps of 0.2 ms were generated for the two conditions, respectively. The diffraction-limited laser spot is defined by a normalized Gaussian intensity profile with a full width at half maximum (FWHM) of 0.6 μm, and maximal value of 1. A molecule which reaches the ROI containing the virtual laser spot, is counted with an intensity equal to the value of this Gaussian function, at the location *r* from the center of the laser spot. The resulting intensity fluctuations over time were analyzed by computing the autocorrelation function. To mimic the impact of photo-bleaching in the probing area, we introduced in the simulations a small photo-bleaching rate, proportional to the local laser intensity. Hence, the photobleaching rate also follows a Gaussian distribution with a maximal rate *k*_*off*_^*Bleach*^ at the beam center (*k*_*off*_^*Bleach*^ = 0.4 s^−1^, 10 times less than for FRAP). Photobleaching had more impact on the autocorrelation function when calculated for the slower molecules in the adhesive contact. FCS simulations were repeated 10 times, and the corresponding autocorrelation functions normalized to their initial value, were averaged.

### SRI simulations

To mimic STORM experiments that rely on the dense labeling of GFP-Nrx1β bound to Nanobody-Alexa647, a large number of molecules were introduced in the virtual cell (70,000 corresponding to 125 mol/μm²). After the diffusion/trapping steady-state has been reached or imposed, the simulation is paused and all diffusion coefficients are set to zero to mimic cell fixation. This procedure accelerates the calculator which does not have to compute new positions at each time frame and just updates fluorescence states. Alternatively, to take into account the fact that a fraction of transmembrane molecules may still be mobile even after fixation with aldehydes^49^, one can impose slow diffusion coefficients. The switch-on rate *k*_*on*_^*Fluo*^ at which fluorescent dyes spontaneously emit light was determined by measuring the fluorescence intensity collected from single Alexa647-conjugated GFP Nanobody molecules bound to the glass coverslip during a STORM sequence, and counting the number of peaks (mean ± sem = 2.5 ± 0.3 peaks over a time period of 420 sec, n = 20 molecules analyzed, giving *k*_*off*_^*Fluo*^ = 0.006 s^−1^). The switch-off rate *k*_*off*_^*Fluo*^ was determined by taking the inverse of the number of time frames during which single Alexa647-conjugated GFP Nanobodies emitted light before entering again the non-emitting state (5.4 ± 0.5 frames of 20 ms, 87 events analyzed), giving a value of *k*_*off*_^*Fluo*^ = 9.3 s^−1^. The on-off duty cycle δ = *k*_*on*_^*Fluo*^ /(*k*_*on*_^*Fluo*^+ *k*_*off*_^*Fluo*^) is the fraction of time that fluorophores spend in the light-emitting state, and equals here 0.00064, very close to reported values for single Alexa647 dyes in MEA-based STORM buffer ^26^. The number of detected molecules per plane in the field of view was around N = 45, corresponding to a total number N/δ = 70,300 actual molecules in the cell geometry that was imaged. Then, simulations were run for 80,000 frames of 20 ms each (total time of 1,600 sec), and a single 16-bit image was generated which contained the integration of all molecule localizations throughout time. To generate a higher number of detected molecules for CNN applications, the parameter *k_on_^Fluo^* was multiplied by a factor of 5 (to 0.03 s^−1^) to mimic the increase in fluorescence emission induced by the 405 nm laser. Three parameters are used to render the super resolution image: the intensity associated with a single detection; the zoom factor which is the ratio between the pixel sizes of the super-resolved image and the low resolution reference picture (a 5-fold zoom corresponds to a pixel size of 32 nm in the high resolution image); and the localization precision, which corresponds to the standard deviation of the Gaussian distribution used to spread detections around the theoretical position of the molecule (σ = 25 nm, FWHM = 58 nm). The SRI still image is saved as a TIFF file.

### Use of FluoSim to train deep CNNs for fluorescence image reconstruction

#### Deep-STORM training procedure

To assess the ability of FluoSim to train deep learning algorithms, we used Deep-STORM ^19^, a Convolutional Neural Network (CNN) designed to localize the positions of fluorescence emitters from dense labeling microscopy images to produce super-resolved images. Deep-STORM is trained with several thousand image pairs: a low resolution fluorescence picture and its associated super-resolved image. The ImageJ plugin ThunderSTORM ^50^ was previously used to generate hundreds of simulated images (of size 64 × 64 pixels) containing randomly positioned emitters, together with a text file containing their positions ^19^. These simulated images depend on several parameters such as camera specifications, point spread function (PSF) of a single emitter, signal-to-noise ratio, and density of emitters. The images and the localization files together with a scaling factor were processed in the MATLAB^®^ (MathWorks) script provided by the authors (*GenerateTrainingExamples.m*) to format and expand the training set, resulting in several thousand pairs of low- and high-resolution images. Each pair contains a randomly selected cropped image of 26×26 pixels and its high resolution counterpart scaled by a zoom factor (typically 4 or 8). These pairs, exported in a single file, are given to the Python script (*Training.py*) to train the network, which provides as output two files containing the network weights and the mean and standard deviation of each image of the training data set, respectively. The weights can ultimately be provided to the testing Python script (*Testing.py*) to create the Deep-STORM network used to reconstruct super-resolution images. Both Python and MATLAB scripts are freely accessible from the Deep-STORM project web page: https://github.com/EliasNehme/Deep-STORM. Deep-STORM and its dependencies were installed on a 64 bits Ubuntu (18.04.2 LTS) workstation, equipped with an Intel Xeon CPU (E5-1607 @ 3.00 GHz x4) together with 40 GB RAM and a 24 GB memory Nvidia P6000 graphics card.

#### Reconstruction of simulated microtubules

To validate the procedure, we first trained a Deep-STORM network using ThunderSTORM, with simulated microtubule datasets that were generated for a contest to evaluate software packages for single molecule localization microscopy ^51^, and are available on the EPFL website: http://bigwww.epfl.ch/smlm/challenge2013/index.html. We generated with ThunderSTORM 200 images of 64 × 64 pixels containing randomly positioned emitters (density = 0.5 μm^−2^). Each emitter was set to produce a Gaussian diffraction spot in the simulated images, with a standard deviation ranging between 115 nm and 180 nm and a peak intensity ranging from 110 to 440 grey levels. No intensity offset was added but a Poisson noise was applied to each pixel. A zoom factor of 8 was set in the MALTAB script to expand the training dataset producing the pairs of low/high-resolution images used to train the Deep-STORM network. This network was able to reconstruct a high resolution image of the simulated microtubules, as described ^19^.

Another Deep-STORM network was then trained with simulated data generated by FluoSim. A square region of 64 × 64 pixels with a 100 nm pixel size was defined, and populated with 20 emitters to reach an emitter density of ~0.5 μm^−2^. To randomize the positions of the emitters between consecutive frames, the diffusion coefficient of the emitters (*D* = 2 μm²/s) and the simulation time step (*δt* = 1 s) were chosen so that the displacements were of the same magnitude as the region length, i.e. 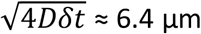. Each emitter was set to produce a Gaussian PSF of 120 nm standard deviation and of peak intensity equal to 110 grey levels in the recorded frames. Here, no intensity offset was added and no Poisson noise was applied to the simulated images. An SPT simulation of 200 steps was performed with the Stack Export option enabled. The resulting image stack and localization file were then processed by the MATLAB script to produce 5,000 pairs of low/high resolution images (Zoom Factor = 8), which were used to train the Deep-STORM network. The resulting network was tested on the simulated microtubules and compared to the ground truth and the image reconstructed with the ThunderSTORM-based network.

#### Reconstruction of real microtubules and Nrx1β-Nlg1 cell-cell contact

To reconstruct our own STORM experiments using Deep-STORM networks, we had to generate specific training sets which reproduce our experimental conditions. Two networks were educated, one trained with ThunderSTORM and one with FluoSim. In both cases 200 square images of 64×64 pixels at pixel size = 160 nm were produced. Each frame contained 20 emitters resulting in an emitter density of 0.5 μm^−2^; for the FluoSim simulation, we followed the same procedure to randomize the positions as for the simulated microtubules. Each emitter was set to produce a Gaussian PSF with standard deviation ranging from 170 nm to 212 nm in the ThunderSTORM simulation, and with a standard deviation equal to 192 nm in FluoSim. In both situations, the PSF maximum was set to 110 grey levels, an intensity offset equal to 30 was added to the images, and a Poisson noise was applied so that each resulting pixel value was randomly taken in a Poisson probability distribution of mean (and hence variance) equal to the sum of the raw pixel intensity value and the intensity offset. The generated low resolution image stacks and localization files were processed by the MATLAB script to produce 5000 pairs of low/high resolution images (Zoom Factor = 4). The Zoom Factor was set to 4 to limit the amount of GPU memory needed to reconstruct the large STORM experiment images (118 × 284 pixels for the Real MT and 168×288 pixels for the Cell Cell contact). Two Deep-STORM networks were finally trained using the data originating from both ThunderSTORM and FluoSim simulations, and were used to reconstruct real microtubules and the Nrx1β-Nlg1 cell-cell contact from STORM experiments in both low and high emitters density situations. STORM experiments in the low density regime resulted in a large amount of images (~50,000) which could not be processed directly with the provided Python script, because of saturation in computer memory. To overcome this problem, the STORM images were reconstructed by batches of 200 images, each batch giving a single super-resolved image. Once reconstructed, the single super-resolved images were summed up to produce the final STORM image.

## Acknowledgements

We thank M. Missler and P. Scheiffele for the generous gift of plasmids; B. Tessier, Z. Karatas, and S. Benquet for molecular biology; R. Sterling and the cell culture facility of the Institute; J.B. Sibarita, C. Butler, and A. Kechkar for the PALM-Tracer program, F. Levet and L. Barusseau for programming tips, E. Nehme for useful advice on Deep-STORM, M. Sainlos for the gift of dye-conjugated GFP nanobody, O. Rossier and G. Giannone for scientific insight, and A. Mehidi and T. Orré for useful feedback.

This work received funding from the Fondation pour la Recherche Médicale (“Equipe FRM” DEQ20160334916), French Ministry of Research, Agence Nationale de la Recherche (grants « SynAdh » ANR-13-BSV4-0005-01 and « Synthesyn » ANR-17-CE16-0028-01), the national infrastructure France BioImaging (grant ANR-10INBS-04-01), and the Conseil Régional Aquitaine (« SiMoDyn »).

## Author contributions

M.L. and O.T. designed FluoSim, performed the FCS experiments, and wrote the user manual. M.L. developed FluoSim and carried out the simulations. M.L. and I.C. performed the SPT and FRAP experiments, and made the figures. I.C. carried out the dSTORM experiments. E.B. and M.N. installed and run the CNN Deep-STORM. O.T. came up with the original idea, supervised the work, and wrote the manuscript. All authors reviewed the manuscript.

## Data availability statement

The authors declare that all data supporting the findings of this study are available within the paper and its supplementary information files.

## Supplementary material

**Supplementary Table 1.** Simulator parameters.

| *Category* | *Parameter* | *Notation* | *Unit/format* |
| --- | --- | --- | --- |
| *Geometry* | Reference image |  | .TIFF, PNG, JPG, GIF |
| | Image pixel size | | Pixels/$\mu\text{m}$ |
|  | Region file |  | .rgn or .roi |
| | FRAP laser spot | | 2.8 $\mu\text{m}$ |
| | FCS laser spot width | $\sigma$ | 0.25 $\mu\text{m}$ |
| *Molecules* | Number* |  | 1-150,000 |
|  | Initial position |  | Random or estimated |
| *Times* | Length scale of simulations* |  | 1,000-100,000 frames (40-1600 s) |
| | Time step | $\Delta t$ | 2-100 ms |
| *Diffusion coefficients* | Outside contact | $D_{out}$ | 0.3 $\mu\text{m}^2/\text{s}$ |
| | Inside contact | $D_{in}$ | 0.3 $\mu\text{m}^2/\text{s}$ |
| | Trapped | $D_{trap}$ | 0.04 $\mu\text{m}^2/\text{s}$ |
| | Crossing probability | $P_{crossing}$ | 25-70% |
| *Kinetics* | Binding rate | $k_{on}$ | 0.15 $\text{s}^{-1}$ |
| | Unbinding rate | $k_{off}$ | 0.015 $\text{s}^{-1}$ |
| *Photophysics* | Switch-on rate* | $k_{on}^{Fluo}$ | 0.005-5 $\text{s}^{-1}$ |
| | Switch-off rate* | $k_{off}^{Fluo}$ | 0-10 $\text{s}^{-1}$ |
| | Bleaching rate* | $k_{off}^{Bleach}$ | 0.4-4 $\text{s}^{-1}$ |
| *Export files* | SPT or SRI images |  | .TIFF or multi-TIFF |
|  | SPT trajectories |  | .trc |
|  | SPT histogram |  | .txt |
|  | FRAP curves |  | .txt |
|  | FCS fluctuations and autocorrelation |  | .txt |

**Supplementary Table 2.** Comparison of FluoSim with other packages.

| **Software** | **2D/3D** | **Modalities** | **Real-time** | **Fluorescence** | **Reference** |
| --- | --- | --- | --- | --- | --- |
| *FluoSim* | 2D | SPT, FRAP, PAF, FCS, PALM, STORM, uPAINT | Yes | Yes | This study |
| *MCell* | 3D | Diffusion, multi-state reactions | No | No | 12,17 |
| *Meredys* | 3D | Diffusion, multi-state reactions | No | No | 18 |
| *SRSim* | 3D | Diffusion, multi-state reactions | No | No | 16 |
| *FERNET* | 3D | FCS | No | Yes | 15 |
| *pyFRAP* | 3D | FRAP | No | Yes | 13 |
| *SuReSim* | 3D | SRI | No | Yes | 14 |

## Supplementary Software

### Supplementary Software

Upon request to the corresponding author, the FluoSim software can be provided as a compressed.zip file to be installed on a computer equipped with Windows operating system. An accompanying user manual describes step-by-step how to use the software. It includes examples of simulations that were used to fit the experimental data contained in the manuscript.

### Supplementary Movie: FluoSim demonstration

To visualize a high definition movie demonstrating the various imaging modalities of FluoSim, please go to the following website: http://www.iins.u-bordeaux.fr/SOFTWARE-285?lang=en

## Supplementary Figures 1-4

**Figure S1.**
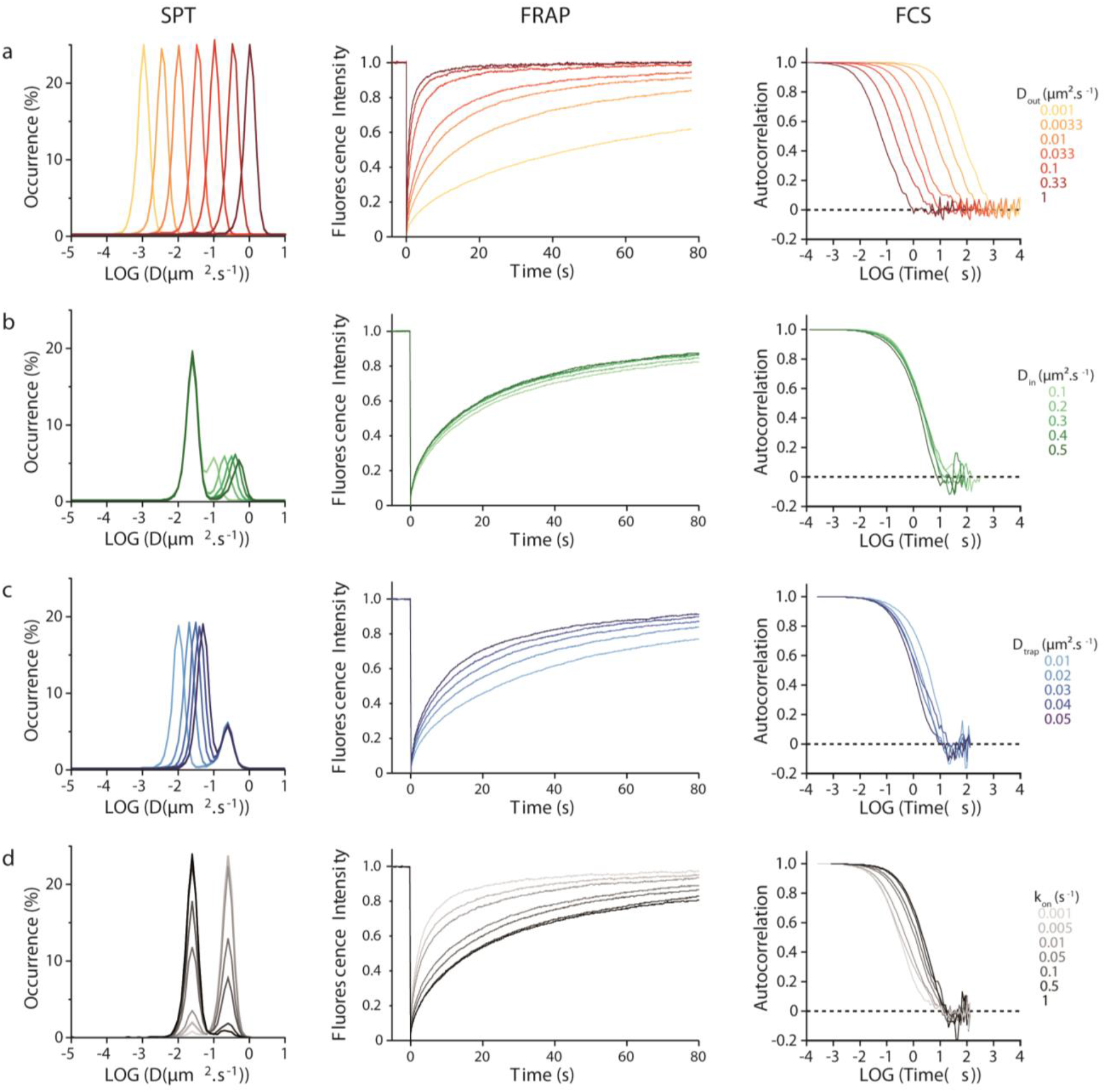
Effect of model parameters on SPT, FRAP and FCS simulations. **(a**) Effect of varying *D*_*out*_ (from 0.001 to 1 μm².s^−1^) on SPT, FRAP and FCS simulations outside the adhesive contact. Note the right shift of the distribution of diffusion coefficients in SPT, the increase in fluorescence recovery for FRAP, and the left shift of the autocorrelation function in FCS, as *D*_*out*_ increases. **(b**) Effect of varying *D*_*in*_ (from 0.1 to 0.5 μm².s^−1^) on the SPT, FRAP and FCS simulations inside the adhesive contact, the other parameters being held constant (*D*_*out*_ = 0.25 μm².s^−1^, *D_trap_* = 0.025 μm².s^−1^, *k*_*on*_ = 0.15 s^−1^, *k*_*off*_ = 0.015 s^−1^). Note the modest effect of *D*_*in*_ on the FRAP and FCS types of curves, which is due to the fact that only a small proportion of molecules diffuse freely at Din within the contact (i.e. most molecules are trapped, due to the high ratio *k*_*on*_/*k*_*off*_ = 10). In SPT, the small peak of diffusing molecules gradually shifts to the left as *D*_*in*_ decreases from *D*_*out*_ to *D_trap_*. **(c)** Effect of varying *D_trap_* (from 0.01 to 0.05 μm².s^−1^) on SPT, FRAP, and FCS simulations inside the adhesive contact (*D*_*out*_ = *D*_*in*_ = 0.25 μm².s^−1^, *k*_*on*_ = 0.15 s^−1^, *k*_*off*_ = 0.015 s^−1^). Note the left shift of the distribution of diffusion coefficients in SPT, the decrease in fluorescence recovery for FRAP, and the right shift of the autocorrelation function in FCS, as *D_trap_* decreases. **(d)** Effect of varying *k*_*on*_ on SPT, FRAP, and FCS simulations inside the adhesive contact (*D*_*out*_ = Din = 0.25 μm².s^−1^, *D_trap_* = 0.025 μm².s^−1^, *k*_*off*_ = 0.15 s^−1^). Note that in SPT the slowly moving population centered at *D_trap_* increases with *k*_*on*_, while the highly mobile population centered at *D*_*out*_ decreases concomitantly. Increasing *k*_*on*_ also slows down the fluorescence recovery in FRAP, and induces a right shift of the FCS curve.

**Figure S2.**
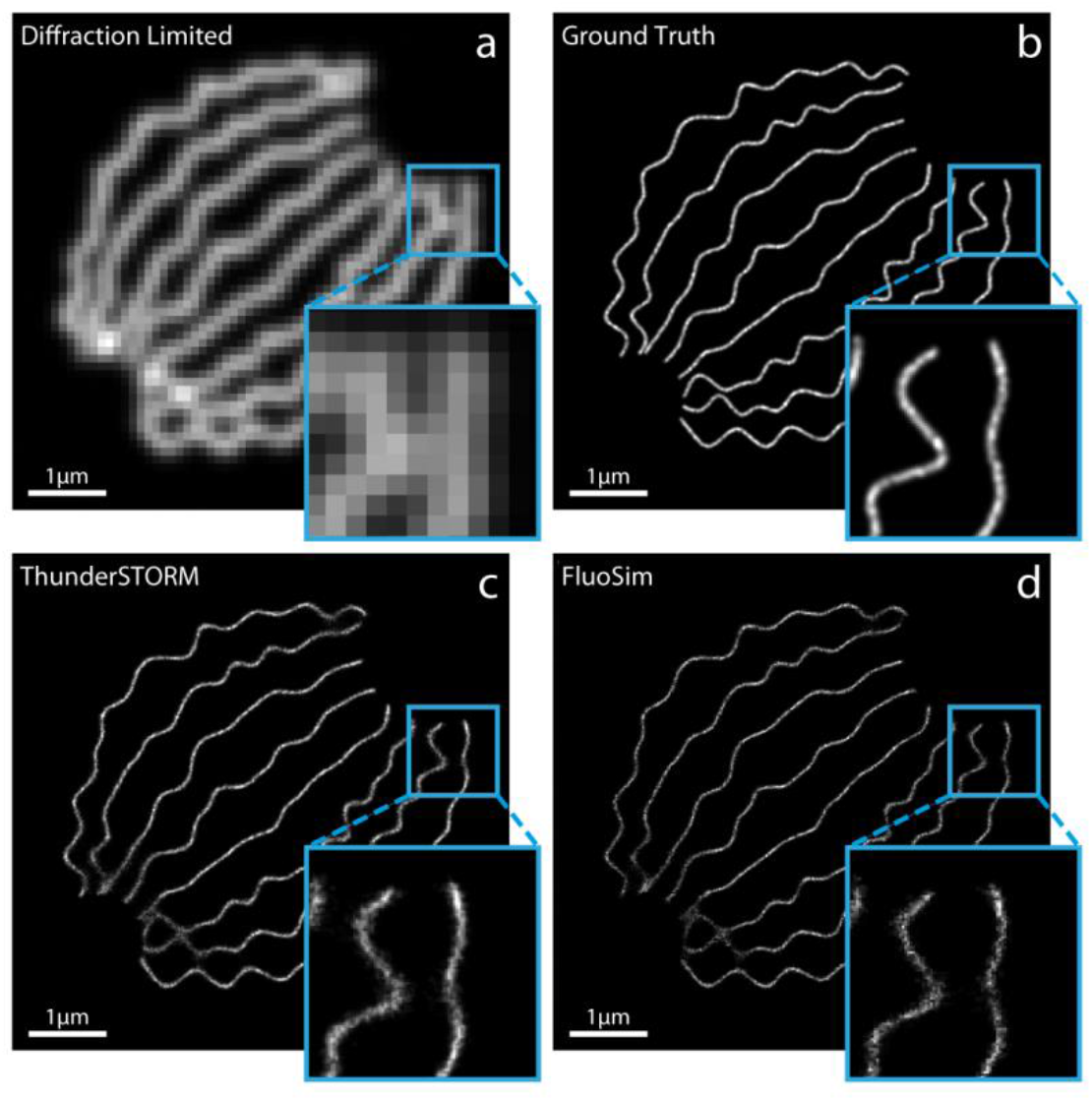
Reconstruction of super-resolved images of simulated microtubules using deep CNN trained with FluoSim. **(a)** Maximal intensity projection image of 350 frames of simulated single molecules randomly placed at high density along virtual microtubules. **(b)** Ground truth image of simulated microtubules taken from the EPFL website ^51^. **(c)** Image reconstructed by the CNN from the single molecule microtubule stack, after training with ThunderSTORM. **(d)** Image reconstructed by the CNN from the single molecule microtubule stack, after training with FluoSim. Note that the images reconstructed by the CNN are close to the ground truth, using either ThunderSTORM or FluoSim training. Insets show zooms on several microtubule ends to highlight the reconstruction accuracy.

**Figure S3.**
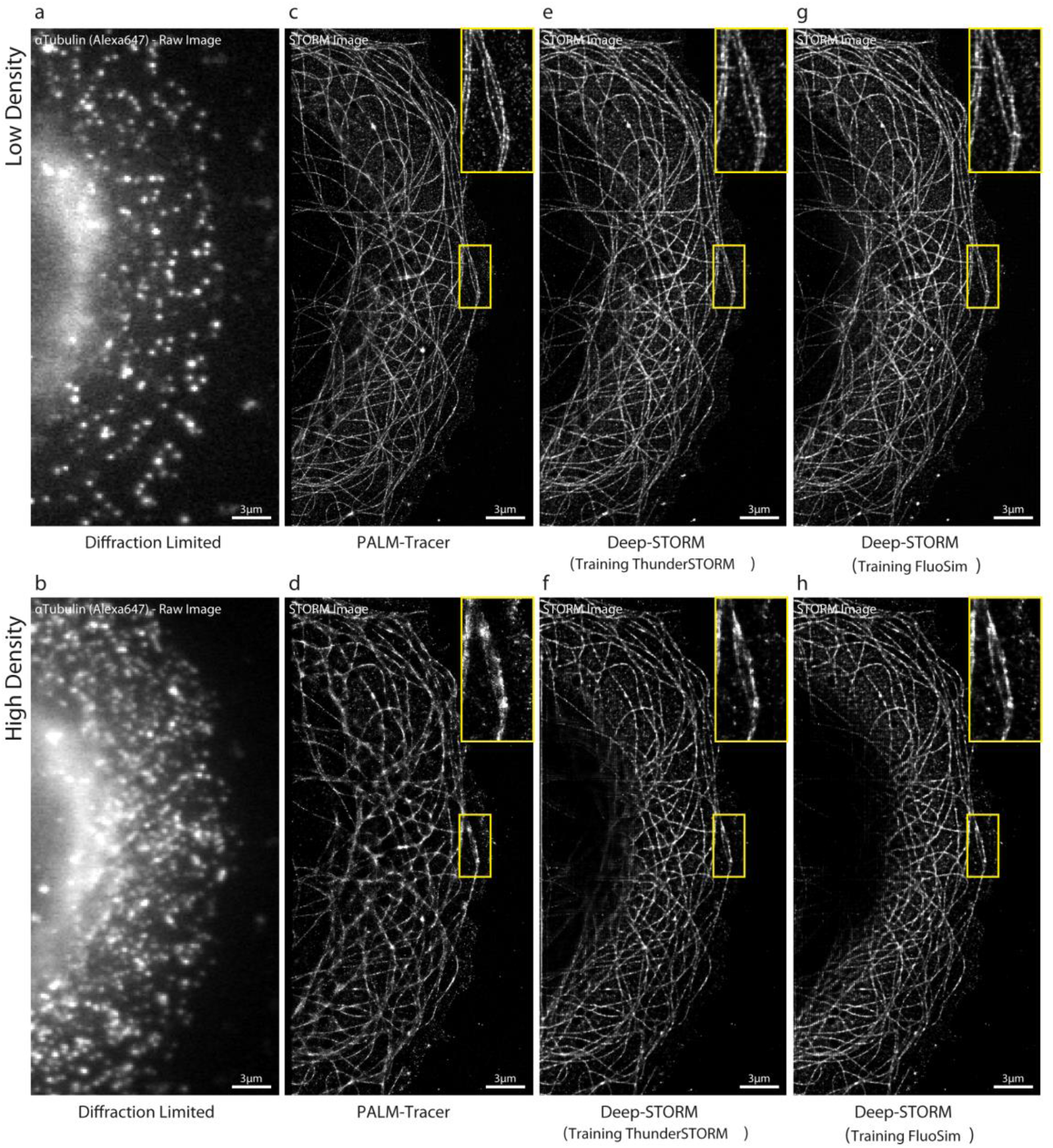
Reconstruction of super-resolved images of real microtubules using deep CNN trained with FluoSim. **(a, b)** Single plane images of microtubules in COS-7 cells, labeled with primary α-tubulin antibody followed by Alexa647-conjugated secondary antibody, and acquired under STORM conditions at low or high density, respectively (obtained for two different values of 405 nm laser power). **(c, d)** Images of microtubules reconstructed using PALM-Tracer from a stack of 48,000 frames at low molecule density, or a stack of 4,000 frames at high molecule density, respectively. Note that the image resolution is degraded at high molecule density because single molecules are too close to one another for proper centroid determination. **(e, f)** Images of microtubules reconstructed by the CNN trained with ThunderSTORM, from low and high molecule density stacks, respectively. **(g, h)** Images of microtubules reconstructed by the CNN trained with FluoSim, from low and high molecule density stacks, respectively. Note that the CNN performs well at both low and high molecule density, thereby offering a significant temporal gain in the image acquisition process.

**Figure S4.**
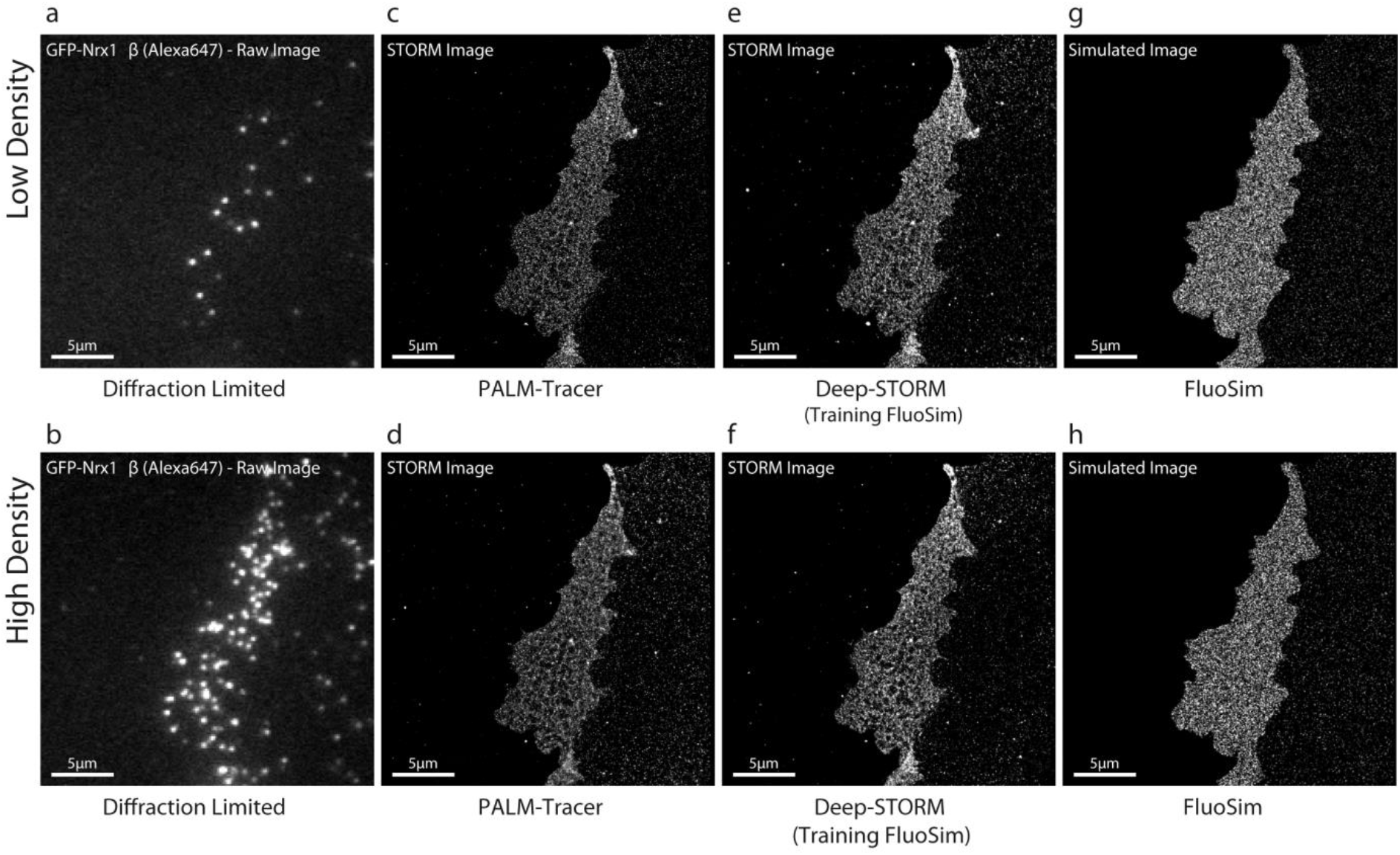
Reconstruction of super-resolved images of Nrx1β-Nlg1 cell contacts using deep CNN trained with FluoSim. **(a, b)** Single plane images of GFP-Nrx1β in COS-7 cells labeled with Alexa647-conjugated GFP nanobody and acquired under STORM conditions, at low or high density, respectively. **(c, d)** Images of Nrx1β-Nlg1 adhesive contacts reconstructed using PALM-Tracer from a stack of 48,000 frames at low molecular density (~30 molecules per frame in the field of view), or a stack of 4,000 frames at high molecular density (~150 molecules per frame in the field of view), respectively. **(e, f)** Images of Nrx1β-Nlg1 contacts reconstructed by the CNN trained with FluoSim, from low and high molecule density stacks, respectively. **(g, h)** SRI images were directly generated by FluoSim at low and high molecule density, respectively. 47,000 molecules were entered in the simulator for a cell surface area of 544 μm², with parameters *k*_*on*_ = 0.15 s^−1^, *k*_*off*_ = 0.015 s^−1^, *P*_*crossing*_ = 0.81, *k*_*on*_^*Fluo*^ = 0.006 s^−1^ (low density) or 0.03 s^−1^ (highdensity), and *k*_*off*_^*Fluo*^ = 9.3 s ^−1^. The total number of detections at low versus high molecular density was 1,436,260 vs 462,607 for experiments and 1,434,610 vs 615,262 for simulations.

